# Extension of bacterial rDNA sequencing to concurrent epigenetic analysis and its application to 16S meta-epigenetics

**DOI:** 10.1101/2022.02.15.480630

**Authors:** Motoi Nishimura, Tomoaki Tanaka, Syota Murata, Akiko Miyabe, Takayuki Ishige, Kenji Kawasaki, Masataka Yokoyama, Satomi Tojo-Nishimura, Kazuyuki Matsushita

**Author notes:** Address correspondence: Motoi Nishimura, Division of Laboratory Medicine, Clinical Genetics and Proteomics, Chiba University Hospital Chiba, Japan.

## Abstract

Although polymerase chain reaction (PCR) amplification and sequencing of the 16S rDNA region has been used in a wide range of scientific fields, it does not provide DNA methylation information. We describe a simple add-on method to investigate 5-methylcytosine residues in the bacterial 16S rDNA region from clinical samples or flora. Single-stranded bacterial DNA after bisulfite conversion was preferentially amplified with multiple displacement amplification (MDA) at pH neutral, and the 16S rDNA region was analyzed using nested bisulfite PCR and sequencing. 16S rDNA bisulfite sequencing can provide clinically important bacterial DNA methylation status concurrently with intact 16S rDNA sequence information. We used this approach to identify novel methylation sites and a methyltransferase (M. MmnI) in *Morganella morganii*. Next, we analyzed bacterial flora from clinical specimens of small amount and identified different methylation motifs among *Enterococcus faecalis* strains. The method developed here, referred to as "add-on" to the conventional 16S rDNA analysis, is the most clinically used bacterial identification genetic test, which provides additional information that could not be obtained with the conventional method. Since the relationship between drug resistance in bacteria and DNA methylation status has been reported, bacterial epigenetic information would be useful in clinical testing as well. Our analysis suggests that M. MmnI has a promotive effect on erythromycin resistance. 16S rDNA bisulfite PCR and sequencing coupled with MDA at pH neutral is a useful add-on tool for analyzing 16S meta-epigenetics.

## Introduction

Bacterial 16S rDNA (rRNA) gene sequencing has enabled the simple estimation of bacterial species by genetic testing and opened the door to microbiome analysis. Although bacterial 16S rDNA sequencing analyzes a relatively small region, it does not require bacterial culture and can estimate bacterial species with fairly high accuracy. Combined with short-read next-generation sequencing (NGS), 16S rDNA gene amplicon-based metagenomic analysis of bacterial communities is possible and relatively straightforward (1). At present, 16S rDNA sequencing has been widely incorporated in the field of science, including food analysis (2-4), environmental surveys (5, 6), and clinical tests (7-10).

Metagenomic analysis of bacterial DNA has been further developed in recent years and has been combined with epigenomic analysis (11), leading to the discovery of new methylation motifs and novel DNA methylases (12-14). However, this new meta-epigenomic technique is based on sequencing the whole genome and/or all nucleic acids in a sample. Thus, it loses the advantages of 16S rDNA analysis, which analyzes a narrow region of bacterial DNA but can identify a wide range of bacterial genera or species even from small-volume samples (15).

Here, we hypothesize that combining 16S rDNA sequencing with bisulfite polymerase chain reaction (PCR) (16) can add DNA methylation information to 16S rDNA analysis while retaining the existing benefits. Furthermore, since 16S rRNA is has several hairpin structures (17), and a palindrome sequence can form a hairpin structure, 16S rDNA may contain palindromic nucleotide sequences. Prokaryotic DNA methyltransferases recognize palindromic DNA sequences (18), hence 16S rDNA may be a region where DNA methylation is detected. Because the degree of methylation and the number of methylation sites on bacterial DNA differ greatly depending on the species (19), we designed primer sequences for the bisulfite PCR used in this combination analysis, including a degenerated universal primer (Figure 1) targeting the 16S rDNA region that detects 0% to 100% methylcytosine content at the primer site. This universal primer set can target both bisulfite-treated and non-bisulfite-treated bacterial genomic DNA (Figure 1), therefore bisulfite PCR and sequencing can provide information on the methylation status of the 16S rDNA sequence in parallel with the targeted 16S rDNA sequence itself.

**Figure 1.**
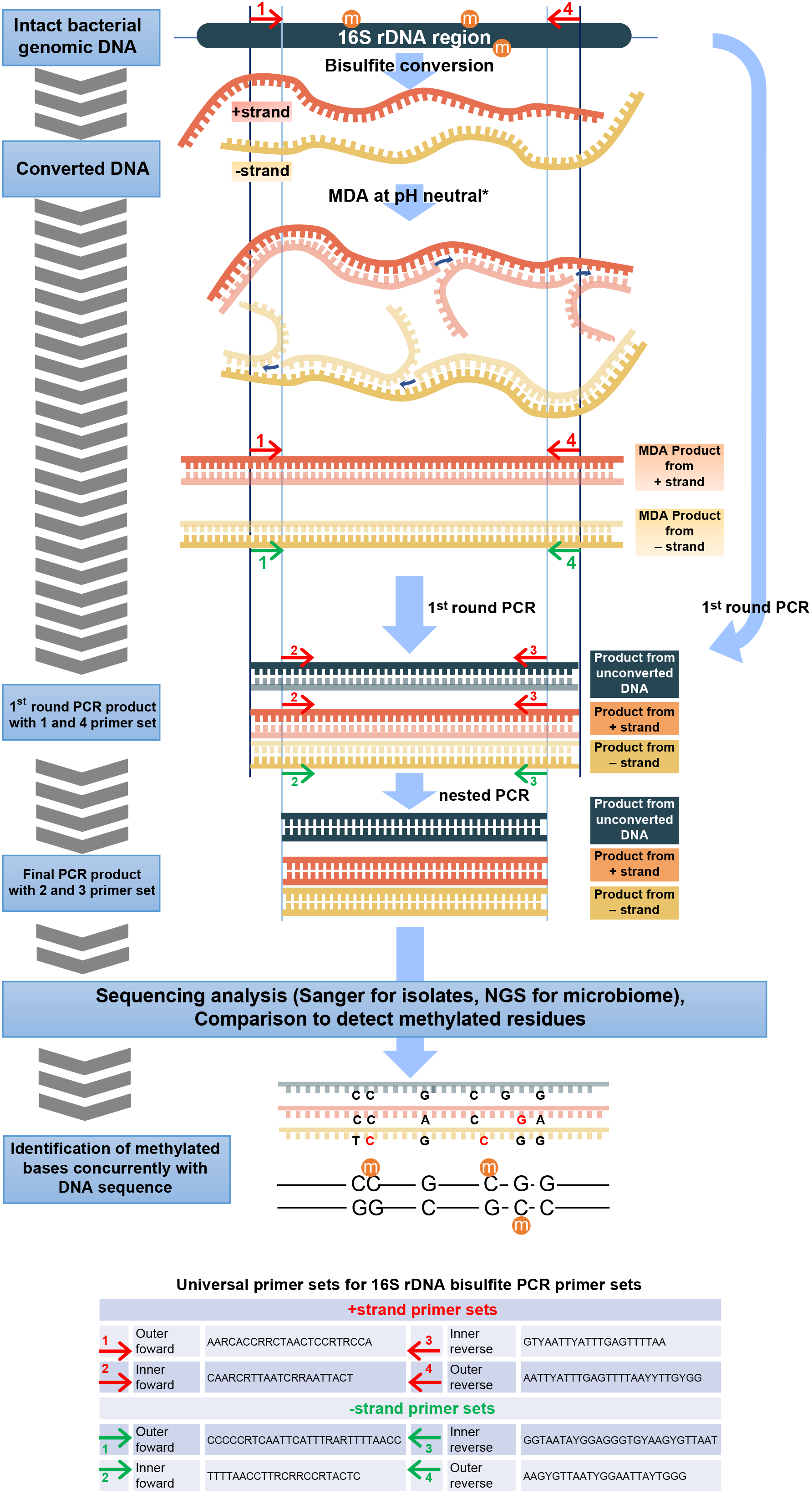
16S rDNA bisulfite PCR and primer sets. The bacterial 16S rDNA V4-V5 region was targeted using bisulfite PCR. Nested PCR primer sets were designed to amplify both intact and bisulfite-converted DNA. The primer sites were chosen to cover several bacterial species and reduce degenerate nucleotides. *MDA at pH neutral was used for the amplification and repair of bisulfite-converted DNA. When amplifying unconverted DNA, the normal MDA method was used. MDA; multiple displacement amplification. R; A or G. Y; C or T. “m” means a methylcytosine base.

## Materials and methods

### Bacterial strains, clinical specimens, and DNA extraction

*Escherichia coli* DNA adenine methyltransferase (dam)+/dcm+ (NEB 5-alpha competent *E. coli*, #C2987) and dam−/dcm− (dam−/dcm− competent *E. coli*, #C2925) strains were purchased from NEB (Ipswich, MA, USA) and cultured in Luria-Bertani (LB) medium. Clinical isolates, namely *K. oxytoca*, *M. morganii*, *P. mirabilis*, *P. aeruginosa*, *S. epidermidis*, *Serratia liquefaciens*, and *E. faecalis*, were cultured on plate agar for routine bacterial identification using MALDI-TOF mass spectrometry (Biotyper, Bruker Daltonics GmbH, Leipzig, Germany) at Chiba University Hospital (20). *Enterococcus faecalis* NBRC strains were purchased from the National Institute of Technology and Evaluation (Tokyo, Japan). Bacterial flora from patient urine samples was washed twice with phosphate-buffered saline and separated via centrifugation at 3,000 × *g* for 5 min at a temperature of 20 °C to 25 °C. Each patient underwent routine clinical bacteriuria tests in which bacterial species were cultured on plate agar and identified using a Biotyper instrument. Bacterial DNA was extracted from either culture medium or separated bacterial flora using an innuPREP Bacteria DNA kit (Analytik Jena GmbH, Jena, Germany) according to the manufacturer’s protocol.

### Bisulfite PCR coupled with MDA at pH neutral

Approximately half of the extracted bacterial DNA was subjected to bisulfite treatment using an innuCONVERT Bisulfite Basic kit (Analytik Jena). The remaining intact DNA was amplified with conventional MDA using a TruePrime WGA Kit (4basebio, Madrid, Spain). The bisulfite-converted DNA was amplified using MDA at neutral pH using the same kit. In MDA at pH neutral, bisulfite-converted single-stranded DNA was amplified untreated with alkaline by mixing the alkaline solution with an acidic solution prior to DNA input. In conventional MDA, intact double-stranded DNA was denatured with alkaline solution before the acid addition and amplification. MDA products from bisulfite-converted or unconverted bacterial DNA were subjected to nested PCR with the primer sets shown in Figure 1. The universal 16S rDNA bisulfite PCR primer sets were designed using the ApE (A plasmid Editor, https://jorgensen.biology.utah.edu/wayned/ape/) and Primer 3 (http://bioinfo.ut.ee/primer3-0.4.0/) software under the following five conditions: i) targeting the V4–V5 region of 16S rDNA; ii) relatively few degenerate nucleotides (R (A, G) or Y (C, T)); iii) primer length > 20 nucleotides; iv) prospects of priming more bacterial species, as expected from the results of BLAST targeting 16S rDNA sites; and v) an expected product size of 364 bp and 377 bp for *E. coli*, which is within the read length of the Ion PGM sequencer.

Bisulfite PCR was performed using the primer sets and nested PCR (Figure 1) utilizing KOD -Multi & Epi-DNA polymerase (TOYOBO, Tokyo, Japan) at a final concentration of 0.02 U/μL. In nested PCR, the final primer concentrations were 0.6 μM in both the first and second PCR steps. The cycling conditions were as follows: 94 °C for 2 min and 15 cycles (first reaction) or 20 cycles (second reaction) at 98 °C for 10 s, Tm °C for 30 s, and 68 °C for 15 s; Tm °C was 56 °C for the positive strand in the first PCR step, 42 °C for the negative strand in the first PCR step, and 52 °C for the second PCR step. Between the two reaction steps, excessive primers were removed with an enzymatic protocol using exonuclease I (NEB) and quick CIP (NEB).

### Sanger and NGS

Sanger sequencing was performed using a 3500xL Genetic Analyzer (Thermo Fisher Scientific, Waltham, MA, USA). A BigDye Terminator v3.1 Cycle Sequencing Kit (Thermo Fisher Scientific) was used for the cycle sequencing reaction. NGS was performed using an Ion Plus Fragment Library Kit, a Hi-Q View OT2 Kit, a Hi-Q View Sequencing Kit, or an Ion 318 Chip Kit v2 (Thermo Fisher Scientific) and a benchtop Ion PGM system according to the manufacturer’s protocol. The DNA fragmentation step was skipped when using the Ion Plus Fragment Library Kit. The Sanger sequencing data and Ion Torrent BAM files were analyzed using CLC Genomics Workbench (Qiagen, CLC bio, Aarhus, Denmark), including running BLAST against the NCBI database (downloaded in May 2021).

### Plasmid preparation

To determine the DNA methyltransferase gene responsible for the consecutive methylation in the *M. morganii* 16S rDNA region (Figure 2B), we selected M.Mom25830ORF6305P and M.Mom25830ORF2065P (designated as M. MmnI in this paper) as candidate genes by searching against a gold-standard dataset in the Restriction Enzyme Database (NEB REBASE) (21). The genes were cloned into the pCold III–Mor1 expression vector between the SacI (Takara Bio) and XbaI (Takara Bio) restriction sites. For the M.Mom25830ORF6305P-expressing plasmid, the gene-specific oligonucleotide primers used were 5′-GGTGAACGGTTCAGACGACT-3′ and 5′-CCTGCGCTACTGTTTCGGTA-3′ in the first round of nested PCR and 5′-ATATGGAGCTCATGAAAAACACTGTTAATTT-3′ and 5′-TACCTATCTAGATCACGTGAAACTTTCAAGACC-3′ in the second round of PCR. For the M.Mom25830ORF2065P-expressing plasmid (designated pCold III-M. MmnI, Figure 3A), the gene-specific oligonucleotide primers used were 5′-TGTTTTTCCGGCCTTCCTGT-3′ and 5′-CATCGGATTTTCAGCCGCTG-3′ in the first round of nested PCR and 5′-CATATGGAGCTCATGATTTTGAAAAAACACCC-3′ and 5′-TACCTATCTAGATTATTTTACCGGCGGTATTG-3′ in the second round of PCR. The entire cloned gene fragments in both plasmids were Sanger-sequenced.

**Figure 2.**
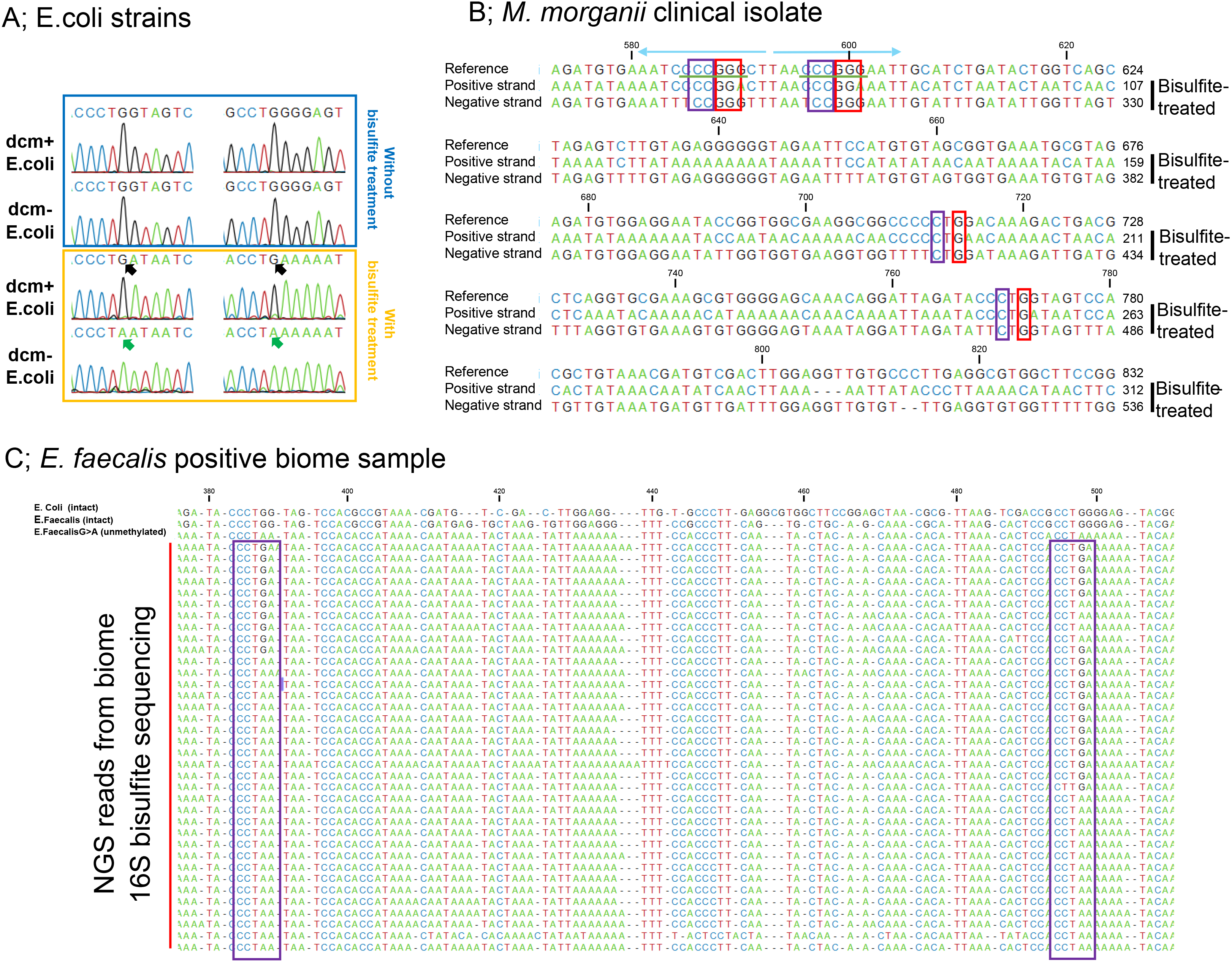
Sequencing the 16S rDNA bisulfite PCR product from clinical isolates and microflora simultaneously provides rDNA sequence information and methylation status. Extracted DNA was treated with bisulfite and amplified with MDA at pH neutral. DNA without bisulfite treatment was amplified using normal MDA. Bisulfite PCR was performed using MDA products as templates, and the resulting PCR products were sequenced. A) Sanger sequencing of bisulfite PCR products from genomic DNA of dcm+ and dcm− *E. coli*. The left side of the figure shows a typical CC(A/T)GG site targeted by dcm methylase and the right half shows another typical CC(A/T)GG site. The difference between dcm+ and dcm− *E. coli* becomes apparent with bisulfite treatment, as shown by the black and green arrows. Green arrows indicate G to A conversion resulting from bisulfite conversion and unmethylated cytosine in the negative-sense strand. The black arrow (not converted) indicates the existence of 5-methylcytosine in the negative-sense strand. B) Summary of bisulfite sequencing from a clinical isolate of *Morganella morganii*. The red boxes indicate 5-methylcytosine in the negative-sense strand. The blue boxes indicate 5-methylcytosine in the positive-sense strand. Note that the four methylated residues at the two SmaI sites (CCCGGG) are underlined in green. The four methylation residues exist in a palindrome-like structure, as indicated by the light blue arrows. The reference sequence was retrieved from NCBI NR_028938.1. C) Implications of diversity in genomic DNA methylation status of *E. faecalis* from a microflora sample. 16S rDNA bisulfite-converted sequencing of MDA products indicated diverse DNA methylation at the CC(A/T)GG site (purple boxes) of the genomic DNA fragment that appears to be from *E. faecalis* in sample number 10_1379 (Supplementary Table S1). In the purple boxes, the letters G written in black in the bisulfite-converted sequences indicate 5-methylcytosine in the negative-sense strand. The reference sequences in the top two columns were retrieved from NCBI NC_000913 (*E. coli*) and NCBI NR_114782.1 (*E. faecalis*) and are the intact sequences, not bisulfite-converted sequences. The reference sequence in the third column from the top is the bisulfite-converted sequence (G>A), assuming that the reference *E. faecalis* sequence does not contain methylated cytosine.

**Figure 3.**
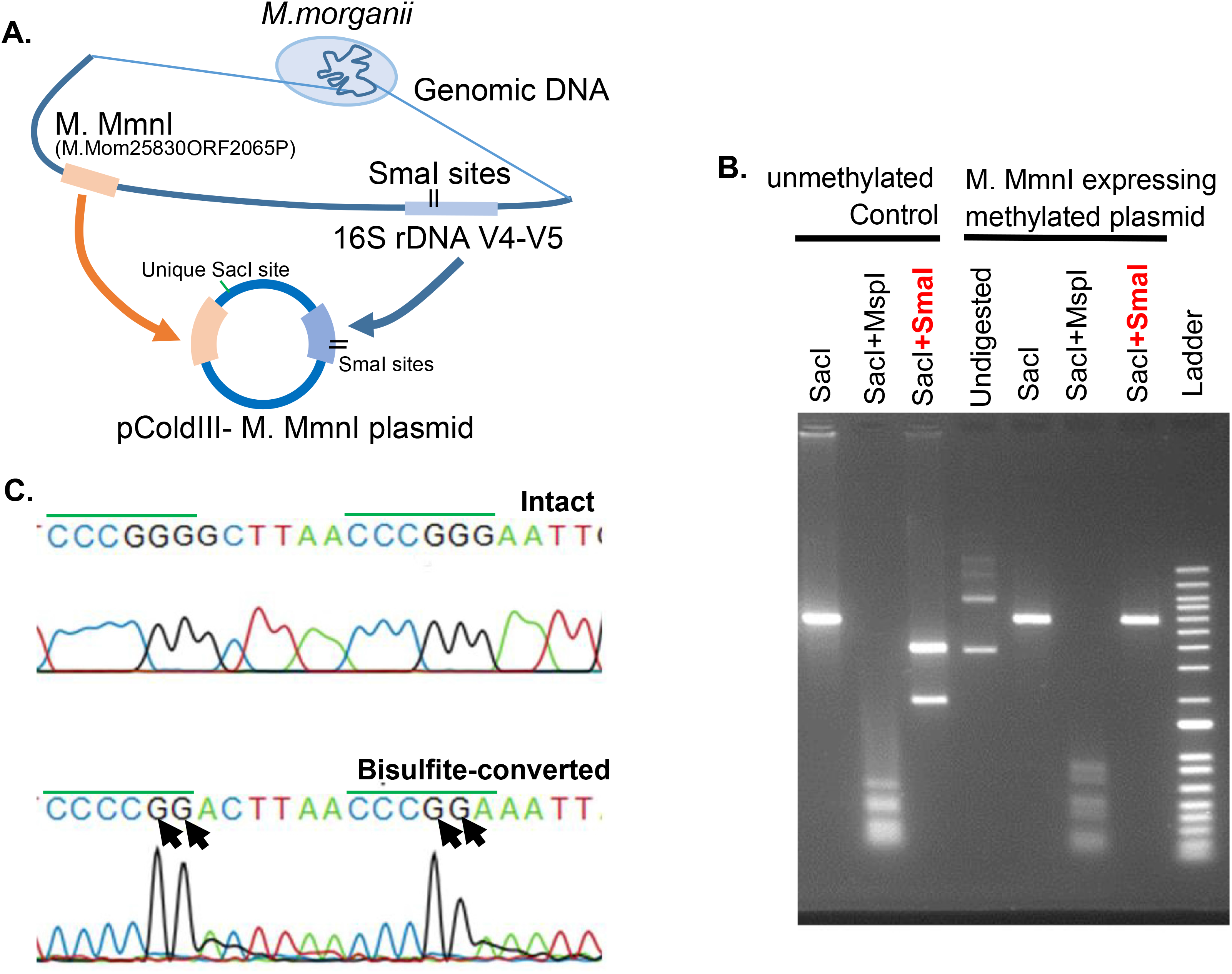
M. MmnI is responsible for four methylated residues at two consecutive SmaI sites within a palindrome-like structure. A) Graphical scheme of pColdIII-M. MmnI (renamed from M.Mom25830ORF2065P) plasmid construction. B) Restriction digestion of the pColdIII-M. MmnI plasmid evaluated via agarose gel electrophoresis. MspI and SmaI were used, where the plasmid contained 37 CCGG and two CCCGGG target sites. Plasmid DNA was linearized using a unique SacI site. Thermo Fisher E-Gel 1 kb plus ladder was employed as a size marker. Control unmethylated DNA was prepared using a Repli-g kit (QIAGEN). C) Bisulfite sequencing of the palindrome-like structure confirmed the presence of four methylated residues in the two consecutive SmaI sites within the structure. The two SmaI sites are overlined in green. Black arrows indicate 5-methylcytosine in the negative-sense strand.

The pCold III –Mor1 expression vector was constructed from pCold III vector (Takara Bio, Kyoto, Japan) at the NgoMIV (NEB) site by inserting the *M. morganii* 16S rDNA V4–V5 region sequence with the following additional NgoMIV site: 5′-GCCGGCCAAGCGTTAATCGGAATTACTGGGCGTAAAGCGCACGCAGGCGGTTGATTGAGTCAGATGTGAAATCCCCGGGCTTAACCCGGGAATTGCATCTGATACTGGTCAGCTAGAGTCTTGTAGAGGGGGGTAGAATTCCATGTGTAGCGGTGAAATGCGTAGAGATGTGGAGGAATACCGGTGGCGAAGGCGGCCCCCTGGACAAAGACTGACGCTCAGGTGCGAAAGCGTGGGGAGCAAACAGGATTAGATACCCTGGTAGTCCACGCTGTAAACGATGTCGACTTGGAGGTTGTGCCCTTGAGGCGTGGCTTCCGGAGCTAACGCGTTAAGTCGACCGCCTGGGGAGTACGGCCGCAAGGTTAAAACTCAAATGAATTGCCGGC-3′.

### Experimental verification of DNA methyltransferase activity

To verify the function of the M. MmnI gene, the plasmids were transformed into *E. coli* dam+/dcm+ (NEB 5-alpha competent *E. coli,* #C2987) and dam−/ dcm− (dam−/dcm− competent *E. coli*, #C2925) strains. The *E. coli* strains were cultured in LB broth (Thermo Fisher Scientific) supplemented with 100 mg/L ampicillin (FUJIFILM Wako Pure Chemical Corporation, Osaka, Japan). Plasmid DNA was isolated using the FastGene Xpress Plasmid PLUS Kit (Nippon Genetics, Tokyo, Japan). SacI was used to linearize the pCold III-M. MmnI plasmid DNA, which was then inactivated by heat at 70 °C. The methylation status was assayed via enzymatic digestion using DNA-methylation-sensitive SmaI (NEB) and -insensitive MspI (NEB) restriction enzymes (Figure 3B). Unmethylated pCold III-M. MmnI plasmid DNA was prepared by amplification using a Repli-g mini kit (Qiagen). We further verified the methylated motifs of M. MmnI. Briefly, half of the pCold III-M. MmnI plasmid DNA extracted from the transformed *E. coli* dam−/dcm− cells was subjected to bisulfite conversion. Both converted and unconverted DNA were subjected to restriction digestion followed by agarose gel electrophoresis (Figure 3B) and to Sanger sequencing as described above (Figure 3C).

### Statistical analysis

All statistical analyses, including calculation of standard deviations, were performed using Excel 2016 (Microsoft, Redmond, WA, USA) with the add-in software Statcel 3 (OMS Publishing Inc., Saitama, Japan).

### Ethics

The present study design, including the associated consent forms and procedures (according to the Ethics Guidelines for Medical Research for Humans in Japan), was approved by the Human Ethics Committee of Chiba University (No. 685). Urine samples were obtained from patients who provided written informed consent, and the samples were then irreversibly anonymized according to the requirements of the ethics committee.

## Results

There are difficulties in using bisulfite PCR on denatured DNA with bisulfite treatment and in sequencing the PCR products during 16S rDNA analysis. One reason for this is that the target DNA is denatured, becomes single-stranded by bisulfite treatment, and is damaged. Bisulfite treatment for methylcytosine-selective hydrolysis causes a significant level of target degradation (22). Even if the target DNA is not damaged, double-stranded DNA (dsDNA) is often more easily amplified than single-stranded DNA (ssDNA) (Supplementary Figure S1). Thus, in 16S rDNA bisulfite PCR, treated target ssDNA is at risk of being less amplified than non-denatured contaminated dsDNA. Further, the possibility of bacterial contamination from the lab environment cannot be completely ignored, especially because contaminating bacteria naturally has double-stranded 16S rDNA.

To reduce the contamination risk and repair degraded DNA and taking advantage of the fact that ssDNA can be preferentially amplified using neutral conditions via multiple displacement amplification (MDA at pH neutral, Supplementary Figure S1), which is usually performed using DNA treated in alkaline conditions, we performed bisulfite PCR of the 16S rDNA region.

Meta-epigenomics analysis assumes that when methylation analysis is randomly performed on DNA molecules in a sample, each sequence is not necessarily a sequence that can directly infer the species from which it originates. Even before meta-epigenomics, there are examples of epigenomic studies of isolated and cultured bacteria (23,24). Whole genomic bisulfite-treated genomic DNA of the *E. coli* K-12 strain has been sequenced (23). A recently published epigenomics method using methylated-cytosine selective restriction enzymes (24), unlike previous meta-epigenomic analyses (11), does not require a single-molecule sequencer and does not change the large-scale analysis of non-targeted, random DNA molecules.

Because the sequence information of the 16S rDNA region is powerful for identifying bacterial species, this analysis has been widely adopted in clinical tests and biome analysis. If DNA methylation analysis can be performed completely in parallel with conventional 16S rDNA analysis, then DNA methylation information can be obtained by targeting regions that contain information that can more efficiently lead to the identification of bacterial species.

Here, we report that 16S rDNA bisulfite PCR coupled with MDA at pH neutral enables 16S meta-epigenetic analysis by extending 16S rDNA sequencing to concurrent epigenetic analysis, which enables the discovery of novel methylation motifs and DNA methylases.

Although Chiba University Hospital is a medium-sized university hospital in Japan, its laboratory routinely performs more than 50,000 bacterial identification tests per year. In this study, clinical specimens prior to isolation and culture in bacterial identification tests and cultured isolates were used for 16S rDNA meta-epigenetic analysis. Specifically, the bacterial flora was isolated from urine samples by centrifugation. Next, bacterial DNA was extracted and used for microbiome analysis. Bacterial DNA was also extracted from cultured isolates. 16S rDNA methylation analysis was performed on all samples (Materials and methods).

The target region for bisulfite PCR coupled with MDA at pH neutral was the 16S rDNA V4-V5 region. This is because the V4 region is a target for popular 16S rDNA analysis kits [Illumina HiSeq sequencer (25)] but is also convenient for testing the ability to detect 5-methylcytosine. For example, in *Escherichia coli*, the V4-V5 region contains five DNA cytosine methyltransferase (dcm) methylation sites (CC(A/T)GG), which were calculated from the NCBI NC_000913 genomic sequence. This region was used because the target region size can be designed to fit into the read length of a short-read next-generation sequencer, such as a Thermo Fischer Scientific Ion PGM sequencer (Thermo Fisher Scientific, Waltham, MA, USA).

To increase the specificity of bisulfite PCR for this region, a universal primer set for nested PCR was designed for each of the positive-sense (+) and negative-sense (−) strands (Figure 1), thereby allowing effective 16S rDNA bisulfite PCR. Next, the bisulfite PCR products were analyzed using benchtop NGS (Ion PGM) or Sanger sequencing (Materials and methods).

To perform bisulfite PCR coupled with MDA at pH neutral, we extracted genomic DNA from dcm+ and dcm− *E. coli* strains. Next, DNA was treated with bisulfite and amplified with MDA at pH neutral. Untreated DNA was amplified with normal MDA as a control. The MDA product was digested with the restriction enzymes MluCI (NEB, Ipswich, MA, USA) and Sau3AI (Takara Bio, Kyoto, Japan) to determine whether the bisulfite-treated DNA was amplified. The MluCI recognition site is AATT and the Sau3AI recognition site is GATC. Therefore, in the amplified product using DNA that has undergone bisulfite treatment, which converts unmethylated cytosine to uracil, many of the Sau3AI recognition sites will disappear, whereas the number of MluCI recognition sites will increase. As shown in Supplementary Figure S2A, the MDA product showed little change when digested with Sau3AI and was completely digested with MluCI, compared to the undigested condition. This result indicates that almost all amplified DNA was derived from bisulfite-converted DNA and that amplification from unconverted DNA was negligible.

The MDA products amplified using DNA extracted from a clinical isolate (*Klebsiella oxytoca*) and or using DNA extracted from a clinical microbiome sample (urine-derived bacteria) showed the same trend of being less digestible by Sau3AI and more digestible by MluCI (Supplementary Figure S2B, S2C), suggesting that the converted DNA was amplified similarly to the previously tested *E. coli* genomic DNA.

Next, 16S rDNA bisulfite PCR was performed on the amplified MDA products at pH neutral using a nested primer set (Figure 1) and the resulting products were sequenced. Of the 5 dcm methylation sequences [CC(A/T)GG] in the *E. coli* 16S rDNA V4-V5 region (Figure 2A), the difference in methylation between dcm+ and dcm− strains was detected at all dcm methylation sites. Thus, bisulfite sequencing coupled with MDA at pH neutral was able to detect cytosine methylated by dcm methylase in the target region.

Clinical isolates prepared for antibiotic sensitivity testing were then used as a template for MDA at pH neutral and for testing the downstream product (Supplementary Figure S2B). The MDA product was subjected to 16S rDNA bisulfite PCR and sequencing. In genomic DNA from *K. oxytoca*, methylation of CC(A/T)GG sites was detected (Supplementary Figure S3A), similar to sites in dcm+ *E. coli*, as previously reported (26). Besides *K. oxytoca*, genomic DNA was extracted from clinical isolates of several species, including *Proteus mirabilis*, *Pseudomonas aeruginosa*, and *Staphylococcus epidermidis*, and was subjected to 16S rDNA bisulfite sequencing. Within the target region, methylcytosine was not detected in *P. mirabilis* (Supplementary Figure S3B). We observed partial methylation in *Pseudomonas aeruginosa* and *S. epidermidis* (Supplementary Figure S3C, D). For *Morganella morganii*, we observed methylation at the CC(A/T)GG sequence and four methylated residues at two consecutive SmaI sites that were encompassed by a palindrome-like structure (Figure 2B).

Clinical microbiome DNA was successfully amplified with MDA at pH neutral (Supplementary Figure S2C), and 16S rDNA bisulfite PCR and sequencing were performed on the MDA products. DNA extracted from the bacterial flora in urine samples was used as the clinical microbiome sample. These clinical samples were simultaneously subjected to isolation culture and routine bacterial identification tests. As in the clinical isolate experiment, the results of the 16S rDNA bisulfite sequencing were consistent with those of bacterial identification tests by isolation culture and matrix-assisted laser desorption/ionization-time of flight (MALDI-TOF) mass spectrometry (Supplementary Table S1). These results also indicated that there may be diversity in the methylation of the genomic DNA fragment that appears to be from *Enterococcus faecalis* in microbiome sample number 10_1379 (Figure 2C).

Although constitutive genomic DNA methylation at the GCWGC and CCGG sites of an *E. faecalis* isolate was previously reported (27), methylation at the CC(A/T)GG site, as detected in this study, has not been reported. It is difficult to determine from the data shown in Figure 2C whether the diversity of methylation at the CC(A/T)GG site is due to mixing *E. faecalis* isolates with different genomic DNA methylation, because Figure 2C can be interpreted to reflect that methylation at the CC(A/T)GG site varies from cell to cell, even in the same strain. Additionally, it is possible that genomic DNA fragments from unknown or known different bacterial species with the same sequence as *E. faecalis* were mixed and analyzed in Figure 2C. Therefore, we conducted an experiment using clinical isolates and NBRC bank strains of *E. faecalis* to verify whether there is diversity in stable DNA methylation in *E. faecalis*. Indeed, the genomic DNA methylation at the CC(A/T)GG site in *E. faecalis* was different among isolates and strains (Supplementary Figure S4).

In *M. morganii*, four methylated residues at two consecutive SmaI sites within a palindrome-like structure were detected, suggesting the presence of an uncharacterized DNA cytosine-methyltransferase (Figure 2B). Since DNA methylation within such palindrome-like sequences has functional implications other than the restriction-modification system (28), we searched for the enzyme responsible for the consecutive methylation.

M.Mom25830ORF6305P and M.Mom25830ORF2065P, which are presumed to encode DNA methyltransferases in *M. morganii*, were identified using the REBASE database. Next, these genes were each cloned into pCold III expression plasmids along with the palindrome-like structure. These constructs were expressed in the *E. coli* dcm− strain using a previously described method ((11), Figure 3A). The results showed that in M.Mom25830ORF6305P, digestion of the extracted plasmid with SmaI was possible and methylation within the palindrome-like structure did not occur. In M.Mom25830ORF2065P, digestion with SmaI was blocked, indicating methylation of the palindrome-like structure (Figure 3B). Bisulfite sequencing of the palindrome-like structure confirmed the consecutive methylation (Figure 3C). The product of the M.Mom25830ORF2065P gene was designated as methyltransferase MmnI (M. MmnI).

Among the known bacterial DNA methylation sites in the SmaI site, SpyI methylase (M. SpyI) causes similar consecutive methylation with 5-methylcytosine (29). M. SpyI recognizes CCAGG, CCTGG, CCCGG, and CCGGG sequences, resulting in consecutive methylation at the SmaI site (29). Similarly, in the plasmid encoding M. MmnI, methylation was found at these four sequences, suggesting that M. MmnI recognizes these four sequences, and that there is partial homology between the amino acid sequences of M. MmnI and M. SpyI (Supplementary Figures S5). Based on these results, we concluded that M. MmnI is the enzyme responsible for the four methylated residues at two consecutive SmaI sites within the palindrome-like structure.

Since M. SpyI is associated with erythromycin (EM) resistance (29), we investigated whether M. MmnI is also related to the responsiveness of *M. Morganii* to EM (30). The transformed *E. coli* dcm− strain containing the M. MmnI expression plasmid was estimated to have a slightly higher IC_50_ (50 % growth-inhibitory concentration, approximately 70 mg/L) for EM than that of the *E. coli* dcm− strain with the plasmid lacking the M. MmnI open reading frame (approximately 32 mg/L) (Supplementary Figure S6).

## Discussion

The 16S rDNA bisulfite PCR and sequencing method coupled with MDA at neutral pH presented in this paper allows the combination of DNA methylation analysis and 16S rDNA analysis via a universal bisulfite PCR primer set. Thus, analysis of various bacteria (Figure 2A, B, and Supplementary Figure S3) and bacterial flora can be performed using small-volume clinical samples. In contrast with meta-epigenomics (11), where all non-targeted nucleic acid sequences in a sample are analyzed randomly using a single-molecule sequencer, our method targets 16S rDNA analysis and can be described as “16S rDNA meta-epigenetics”. Although the former method provides comprehensive DNA sequence information, including methylcytosine and methyladenine, it requires large sample sizes, for example, 30 L of lake water (11). In contrast, our method can analyze only 2 mL of urine. Obtaining DNA methylation information in the 16S rDNA region is remarkably efficient in identifying the bacteria, from which the methylation originates. However, our method detects methylcytosine but not methyladenine. Since 16S rDNA meta-epigenetics can simultaneously infer species and identify DNA methylation patterns, our results allow us to infer that *E. faecalis* has various methylation patterns depending on the strain or isolate (Figure 2C).

Sequencing of 16S rDNA has been practically incorporated into clinical testing. In clinical laboratories, 50,000 bacterial isolation, culture, and identification tests are conducted annually at our hospital alone, and countless specimens are discarded after testing worldwide. The method reported here can be added on to the conventional16S rDNA identification test and enables DNA methylation analysis of various bacterial species in these discarded specimens. It is important to note that these bacteria were alive at the time of disposal, which means that if DNA methylation is detected using this method (Figure 2B), then living bacteria can be recovered and prepared for further analysis (Figure 3). As shown in Supplementary Figure S3C and D, a partial methylation pattern was detected in both gram-negative (*P. aeruginosa*) and gram-positive (*S. epidermidis*) bacteria. Partial methylation in bacterial genomes was reported in several epigenetic studies (23,31). These studies reported that partial DNA methylation may have some significance in relation to bacterial growth (23). Such cases are among the targets for future verification and further research.

In addition, we found that M MmnI from *M. morganii* methylates four consecutive SmaI sites in the palindrome-like sequence within the 16S rDNA region (Figures 2B and 3) and that M. MmnI may contribute to EM resistance. *M. morganii* is naturally resistant to EM (30). M. SpyI, a DNA methyltransferase with properties similar to M. MmnI, is associated with EM-resistant *S. pyogenes* strains. However, EM resistance has been primarily explained by other drug resistance genes genetically linked to M. SpyI (29). It was recently reported that knockout of dcm DNA methyltransferase, which recognizes CCAGG and CCTGG sequences, suppresses *E. coli* growth at lower EM concentrations (32). Therefore, DNA methyltransferases that recognize CCAGG and CCTGG sequences may be modifying factors in EM resistance, if not the main factor. The relationship between bacterial responses to antibiotics and DNA methylation status (33) is an important subject for future analysis by 16S rDNA meta-epigenetics.

The 16S rDNA region is not large, but it is a prominent informative region that has been targeted for microbiome analysis. Our 16S rDNA bisulfite PCR and sequencing method will enable the simultaneous detection of (i) bacterial species and (ii) associated DNA methylation patterns directly from small amounts of bacterial flora specimens or specimens after isolation, culture, or drug resistance tests. Further, our method will facilitate the detection of unknown DNA methylation patterns and the identification of the responsible methyltransferases. In clinical laboratories, 16S rDNA bisulfite PCR and sequencing will contribute to the elucidation of the relationship between DNA methylation and clinically valuable information such as drug resistance.

## Acknowledgements

We would like to thank Editage (www.editage.com) for English language editing. This work was supported by JSPS KAKENHI Grant Numbers JP21H02823 and JP18K07408.

